# CGMD: Centrality-Guided Maximum Diversity for Annotation-Efficient Fine-Tuning of Pretrained Cell Segmentation Models

**DOI:** 10.1101/2025.11.04.686267

**Authors:** Eiram Mahera Sheikh, Alaa Tharwat, Constanze Schwan, Wolfram Schenck

**Affiliations:** Department of Engineering and Mathematics, Bielefeld University of Applied Sciences and Arts, Bielefeld, Germany

**Keywords:** Cell instance segmentation, Cell Tracking Challenge, Subset selection, Active learning

## Abstract

Pretrained cell segmentation models have simplified and accelerated microscopy image analysis, but they often perform poorly on new and challenging datasets. Although these models can be adapted to new datasets with only a few annotated images, the effectiveness of fine-tuning depends critically on which images are selected for annotation. To address this, we propose CGMD (Centrality-Guided Maximum Diversity), a novel algorithm that identifies a small set of images that are maximally diverse with respect to each other in the pretrained feature space. We evaluate CGMD under an extremely low annotation budget of just two images per dataset for fine-tuning the pretrained Cellpose Cyto2 model on four different 2D+t datasets from the Cell Tracking Challenge. CGMD consistently outperforms six competitive active learning and subset selection methods and approaches the performance of fully supervised fine-tuning. The results show that centrality-guided maximum diversity subset selection enables stable and annotation-efficient fine-tuning of pretrained cell segmentation models. The code is publicly available at: https://github.com/eiram-mahera/cgmd.

## 1. INTRODUCTION

Accurate cell segmentation is fundamental to quantitative microscopy as it underpins downstream analyses such as cell tracking, morphological quantification, and phenotype characterization. Manual segmentation is slow, requires expert supervision, and does not scale. Classical image-processing pipelines based on thresholding, watershed transforms, or morphological operations often fail in the presence of complex cellular structures, overlap, or modality-specific artifacts. Deep learning models have achieved state-of-the-art accuracy [1], but typically require large and carefully annotated datasets. Acquiring such datasets for each experimental condition remains costly and time-consuming.

In this work, we consider supervised training scenarios in which unlabeled data is readily available but annotation cost strictly limits the number of images that can be labeled. Recent efforts have therefore focused on developing pretrained segmentation models that generalize across cell types and microscopy modalities. Cellpose [2, 3] is one such model trained on heterogeneous microscopy data to learn transferable representations of cellular morphology. While these models provide strong out-of-the-box performance, their accuracy can degrade on new and challenging datasets. Fine-tuning such models using a small number of annotated images can substantially improve their performance. However, the effectiveness of finetuning depends critically on which images are chosen. Identifying a small subset of images that will have the greatest impact on down-stream performance after fine-tuning is therefore a central challenge in achieving efficient model adaptation.

This challenge has been addressed in part by active learning and subset selection strategies, which aim to identify the most informative samples for annotation based on criteria such as uncertainty, diversity, or representativeness [4, 5, 6, 7, 8, 9, 10, 11, 12]. These approaches typically involve multiple rounds of sample selection, annotation, and model retraining. While such iterative schemes can be effective, they are impractical for biomedical image segmentation, where annotating a single image may require minutes to hours of expert work and repeated fine-tuning adds significant computational cost. Moreover, most prior studies evaluate their methods under moderate annotation budgets, often tens or hundreds of labeled images which still represent substantial human effort for instance-level segmentation and do not reflect the extreme low-budget scenarios that many laboratories actually face.

To address these limitations, we propose Centrality-Guided Maximum Diversity (CGMD), a novel algorithm designed to select a maximally diverse subset of samples guided by a fixed notion of global centrality. By selecting samples that are maximally diverse with respect to both each other and the dataset center, CGMD promotes broad coverage of dataset-specific variations while avoiding the selection of similar samples. The approach follows the same high-level goals as diversity and representativeness based active learning and subset selection strategies, but is specifically designed for the practical constraints of biomedical image analysis workflows. We evaluate CGMD in the context of annotation-efficient fine-tuning of pretrained cell instance segmentation model where repeated selection and training cycles are impractical

We compare CGMD against representative methods from three major families of active learning and subset selection strategies based on diversity and representativeness: clustering-based selection [13, 14], geometric coverage approaches [15, 16], and diversity-based sampling techniques [17, 18, 19]. These baselines capture the main algorithmic perspectives in prior work and provide a rigorous benchmark. All methods are evaluated under a fixed annotation budget of two images per dataset, reflecting realistic constraints in biomedical imaging. Experiments are conducted using the pretrained Cellpose Cyto2 model [3] on four 2D+t datasets from the Cell Tracking Challenge (CTC)^1^. Across datasets, CGMD consistently outperforms competing baselines and approaches the performance of fully supervised fine-tuning, demonstrating the effectiveness of centrality-guided diversity maximization for annotation-efficient fine-tuning of cell instance segmentation model.

## 2. METHOD

We propose **CGMD (Centrality-Guided Maximum Diversity)**, a novel algorithm that identifies a small subset of informative samples by maximizing the diversity among them. CGMD can operate on any type of feature representation: raw, handcrafted, or learned, and therefore integrates smoothly with pretrained segmentation models. In this work, we apply CGMD to the problem of annotation-efficient fine-tuning of a pretrained cell segmentation model under an extremely small labeling budget.

### 2.1 Feature Extraction and Overview

Figure 1 summarizes the overall workflow. First, we extract feature representations from all microscopy images in the dataset using the pretrained Cellpose Cyto2 model [3]. For each image, we obtain a 256-dimensional feature vector (also called as *style vector* in [2, 3]) that captures global image characteristics including texture, cellular morphology, and imaging modality. Next, the proposed CGMD algorithm operates in this feature space to identify two images that are maximally diverse with respect to each other and with respect to a central reference image. These selected images are then annotated to obtain ground-truth instance segmentation masks. The annotated images are used to fine-tune the pretrained model, adapting it to the dataset. Finally, the fine-tuned model is used to segment images from the same dataset.

**Fig. 1:**
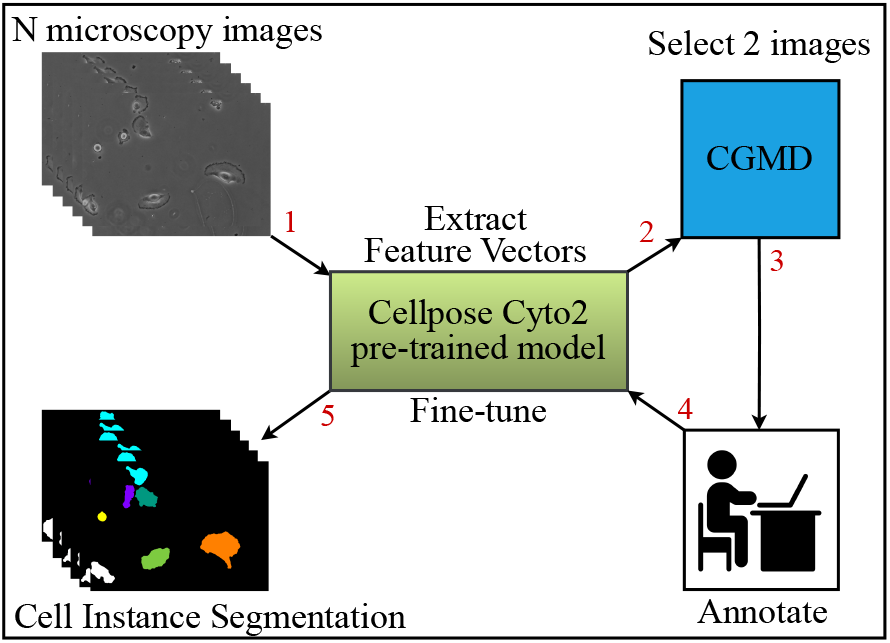
Overview of the proposed CGMD pipeline. Features are extracted from microscopy images using the pretrained Cellpose Cyto2 model. CGMD identifies two images that are maximally diverse and then they are annotated and used to fine-tune the model.

### 2.2 CGMD Algorithm

Algorithm 1 outlines the full procedure. Let 𝒳 = {x_1_, …, x_*N*_} denote the set of *N* feature vectors in ℝ^*d*^, and let *B* denote the annotation budget.

#### Algorithm 1: CGMD

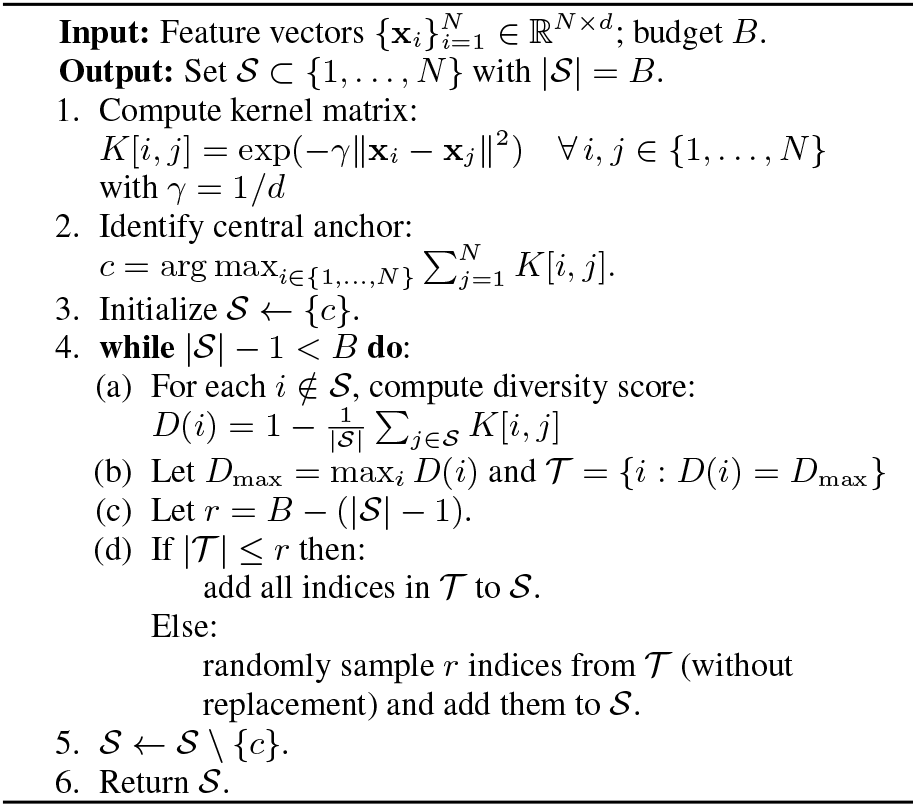

#### 2.2.1 Similarity Computation

We compute a Gaussian radial basis function (RBF) kernel [20] *K* to quantify pairwise similarity between the feature vectors:

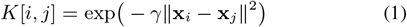

where *γ* controls the rate at which similarity decays with distance between the features. Following a common heuristic in high-dimensional feature spaces, we set:

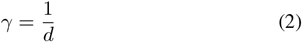

where *d* is the dimensionality of the feature vectors. This scaling maintains a consistent similarity range across datasets with different feature dimensions and provides a stable basis for measuring diversity in the subsequent selection steps. The resulting matrix *K* ∈ ℝ ^*N*^*×*^*N*^ forms the foundation for the CGMD algorithm.

#### 2.2.2 Central reference identification

Next, CGMD identifies a *central reference sample* as the index of the sample whose feature representation has the highest total similarity to all others:

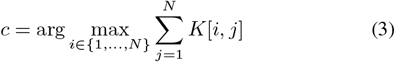

This central reference provides a fixed and deterministic starting point for computing diversity scores. In particular, when the annotation budget is very small, the first reference used in scoring has a strong influence on which samples are ultimately selected. Using a central reference avoids initializing the selection from an arbitrary sample and ensures that diversity is measured consistently across all the samples.

#### 2.2.3 Set initialization

Initialize a set *S* with the *c* obtained from Eq. (3).

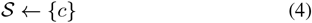

This set is used to store the indices of the samples selected for annotation by the CGMD algorithm. Note, that *c* is never selected for annotation, it is used here because it provides a deterministic starting point for the algorithm.

#### 2.2.4 Diversity scoring and greedy selection

For each feature vector x_*i*_ with index *i* ∉ 𝒮, CGMD computes a *diversity score* that measures how dissimilar x_*i*_ is from the samples already in 𝒮.

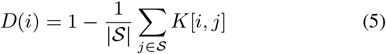

A higher *D*(*i*) indicates that sample *i* is more diverse and less similar to the currently selected subset. At each iteration, the sample (or samples) with the highest diversity score is added to 𝒮. If multiple samples have the same maximum score, CGMD includes all of them if the remaining annotation budget allows; otherwise, it randomly samples from the tied candidates to ensure unbiased tie resolution. This process continues until *B* number of samples are selected.

#### 2.2.5 Central reference removal

After the selection process is complete, the central reference sample *c* is removed from the set 𝒮. This step reflects the distinct role of the central reference in CGMD. The reference sample is introduced solely to guide the computation of diversity scores and to provide a stable and deterministic reference during selection. It is not intended to be part of the annotated subset. In the setting considered here, CGMD is used to fine-tune a pretrained Cellpose Cyto2 model, which already captures generic microscopy image charecteristics. Under such conditions, annotating the reference sample which is, by construction, most similar to all other samples, provides limited additional signal for fine-tuning. Instead, the annotation budget is allocated to samples that achieve high diversity scores relative to this reference and to each other.

### 2.3 Complexity analysis

The computational cost of CGMD is dominated by the construction of the similarity matrix *K*. Computing the RBF kernel for *N* samples of dimensionality *d* requires 𝒪 (*N* ^2^*d*) time and𝒪 (*N* ^2^) memory. The subsequent greedy selection stage adds an additional 𝒪 (*BN*) operations, where *B* is the annotation budget. Since *B*≪*N* in practical scenarios (e.g., *B*=2 in our experiments), the overall runtime is primarily determined by the kernel computation. Consequently, the total complexity of CGMD scales quadratically with the number of samples and linearly with the feature dimensionality. In practice, the algorithm remains efficient due to its vectorized implementation.

## 3. EXPERIMENTS & RESULTS

### 3.1 Datasets

We evaluate CGMD on four 2D+t datasets from the Cell Tracking Challenge. **DIC-C2DH-HeLa** has dense clusters of cells with very weak boundary contrast, DIC-enhanced internal structures, and the coexistence of flat interphase cells with bright and rounded mitotic cells. **Fluo-N2DH-GOWT1** has heterogeneous fluorescence signal inside nuclei due to unlabeled nucleoli and strong variability in intensity across cells. **PhC-C2DH-U373** has halo and shade-off artifacts from phase-contrast imaging combined with highly irregular cell shapes. **BF-C2DL-HSC** has strong visibility of the slide and additional debris that makes cell segmentation by a model difficult. To mitigate this, we crop the images to regions containing cells. In our experiments, we use four slightly different crops to ensure that results are not overly dependent on a specific crop size. Each dataset represents a different modality and cell type with distinct challenges. We briefly summarize the main challenges for each dataset here, for additional details, we refer the reader to the CTC website and papers [21, 22, 23].

### 3.2 Evaluation Metric

We use the **SEG** measure as our main evaluation metric, following the official CTC protocol. It is derived from the Jaccard index:

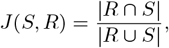

where *S* is a segmented mask and *R* a reference mask. A prediction counts as matching if |*R* ⋂ *S* |*>* 0.5 *R* . Each reference object can be matched at most once. SEG is the mean Jaccard index across all matched objects in a video, yielding values between 0 (no match) and 1 (perfect match). CTC provides two reference annotation sets: sparse *gold truth* (GT) manually drawn by experts, and dense *silver truth* (ST) obtained by fusing outputs from top-performing algorithms. We evaluate against ST masks using the CTC evaluation software.

### 3.3 Baseline Methods

We benchmark CGMD against six competitive active learning and subset selection strategies spanning three methodological families:

#### Clustering-based methods

Clustering provides a natural strategy for subset selection by grouping similar samples and choosing representatives from each cluster. **K-means** clustering [13] partitions data into *k* clusters and selects points closest to centroids ensuring representative coverage of the data distribution. **TypiClust** [14] improves on this idea by choosing typical (high-density) samples within clusters which has been shown to be especially effective in low-budget regimes.

#### Coverage-based methods

Another family of approaches aims to cover the geometry of the data distribution in feature space. **Core-Set** [15] formulates subset selection as a *k*-center problem, where samples are chosen to minimize the maximum distance between any data point and the selected set. **Facility Location** [16] instead casts the problem as submodular optimization, selecting a subset that ensures each point in the dataset is well-represented by at least one chosen element. These approaches balances representativeness and diversity by emphasizing global coverage.

#### Diversity-based methods

These approaches directly optimize for diversity without relying on clustering or explicit geometric coverage. **FADS** [19] is a greedy diversity subsampling algorithm that iteratively selects samples with maximal dissimilarity, producing subsets that capture the variability of large datasets. Determinantal Point Processes (**DPP**) [17, 18, 24] provides a probabilistic formulation, assigning higher probability to subsets with low redundancy. By modeling negative correlations between items, DPPs offer a principled framework for diversity-driven selection.

### 3.4 Experimental Setup

All experiments use the pretrained Cellpose Cyto2 model [3] for feature extraction and fine-tuning. Following common practice, we use sequence 01 for training (subset selection and fine-tuning) and sequence 02 for evaluation across all datasets. Each method selects exactly two images (*B* = 2) for annotation. Models are fine-tuned on the selected images for up to 200 epochs. The optimal number of epochs is chosen by leave-one-out cross-validation (LOOCV) over the two annotated samples, performed independently for each method and dataset. In the full-data regime, we fine-tune on all training images with a validation split of 20% to select the best epoch (max 200). We use the same hyperparameters for all experiments: stochastic gradient descent with a learning rate 0.1 and weight decay 1 *×*10^−4^. To account for stochasticity in subset selection and fine-tuning, every experiment is repeated 100 times. We report mean*±* standard deviation. We also report the performance of the pretrained Cellpose Cyto2 model (without fine-tuning) for context.

## 4. RESULTS AND DISCUSSION

Table 1 summarizes the performance of all methods across the four CTC datasets. Several consistent trends can be observed.

**Table 1:** SEG scores (mean *±* std over 100 runs) for all methods across four 2D+t CTC datasets. Pretrained: Cellpose Cyto2 without fine-tuning. Fully Supervised: fine-tuning using all the images. Boldface indicates the best method using 2 annotated images for fine-tuning.

| Method | BF-C2DL-HSC | DIC-C2DH-HeLa | Fluo-N2DH-GOWT1 | PhC-C2DH-U373 |
| --- | --- | --- | --- | --- |
| Pretrained | 0.763 | 0.020 | 0.906 | 0.258 |
| CGMD (ours) | **0.858 ± 0.008** | **0.842 ± 0.000** | **0.949 ± 0.000** | **0.857 ± 0.000** |
| Core-Set [15] | 0.588 ± 0.275 | 0.807 ± 0.021 | 0.947 ± 0.002 | 0.839 ± 0.021 |
| DPP [17, 18, 24] | 0.615 ± 0.296 | 0.802 ± 0.024 | 0.946 ± 0.003 | 0.844 ± 0.010 |
| FADS [19] | 0.632 ± 0.320 | 0.807 ± 0.021 | 0.944 ± 0.004 | 0.848 ± 0.004 |
| Facility Location [16] | 0.703 ± 0.217 | 0.833 ± 0.000 | 0.946 ± 0.000 | 0.845 ± 0.000 |
| K-means Clustering [13] | 0.723 ± 0.089 | 0.816 ± 0.017 | 0.946 ± 0.001 | 0.844 ± 0.010 |
| TypiClust [14] | 0.699 ± 0.183 | 0.824 ± 0.000 | 0.947 ± 0.000 | 0.842 ± 0.000 |
| Random Sampling | 0.487 ± 0.279 | 0.797 ± 0.040 | 0.942 ± 0.005 | 0.845 ± 0.019 |
| Fully Supervised | 0.902 ± 0.021 | 0.842 ± 0.008 | 0.951 ± 0.001 | 0.850 ± 0.010 |

### CGMD achieves the best overall performance

Across all datasets, CGMD attains the highest mean SEG score. On *BF-C2DL-HSC*, where strong brightfield artifacts and debris make segmentation particularly challenging, CGMD reaches 0.858, substantially higher than the next best method (0.723 for K-means) and far above random sampling (0.487). On *DIC-C2DH-HeLa*, which features densely packed cells with varying morphologies, CGMD achieves 0.842, matching the fully supervised result and outperforming all baselines. For the fluorescence dataset *Fluo-N2DH-GOWT1*, CGMD again yields the top score (0.949), though with smaller margins since most methods perform well for this dataset. Finally, on *PhC-C2DH-U373*, characterized by halo and shade-off artifacts, CGMD achieves 0.857, clearly ahead of competing approaches.

### Performance approaches full supervision

In three of the four datasets, CGMD nearly matches the performance of fully supervised fine-tuning: 0.842 vs. 0.842 on HeLa, 0.949 vs. 0.951 on GOWT1, and 0.857 vs. 0.850 on U373. The only noticeable gap occurs on HSC (0.858 vs. 0.902), likely due to the fact that the video contains over 1,700 frames, and just two images cannot fully represent the dataset. Nevertheless, these results demonstrate that even minimal annotation can yield significant and stable performance gains, providing a strong foundation for incremental extension when more labels become available.

### Baselines exhibit redundancy and instability

Clustering- and coverage-based methods such as Core-Set, Facility Location, and K-means perform reasonably on some datasets but lack consistency across modalities. Their results show high variance, particularly on HSC, where the selected samples often overlap in appearance or fail to capture key variations. Diversity-only methods such as DPP and FADS perform less reliably, sometimes selecting highly distinct but uninformative samples, which leads to unstable fine-tuning outcomes. Random sampling, as expected, remains the weakest and most inconsistent baseline.

### Why CGMD succeeds

CGMD is motivated by a limitation shared by existing strategies: they do not explicitly enforce global representativeness prior to optimizing diversity. K-means selects samples near cluster centers, yielding local rather than global representativeness, while TypiClust emphasizes dense regions and often overlook rare but informative modes. Core-Set and Facility Location prioritize geometric coverage, which at very small budgets biases selection toward boundary points. DPP and FADS maximize mutual dissimilarity, but in the absence of a stabilizing reference can overemphasize weakly informative samples. CGMD overcomes these limitations by conditioning diversity on a fixed notion of global centrality. Rather than favoring locally typical or boundary samples, it prioritizes samples that are maximally distinct relative to the dataset’s dominant structure, thereby reducing redundancy and outlier bias under small annotation budgets.

## 5. CONCLUSION AND FUTURE WORK

We introduced Centrality-Guided Maximum Diversity (CGMD), a novel algorithm that leverages a fixed notion of global centrality to guide diversity-driven sample selection. We demonstrated its effectiveness for selecting the most informative images for fine-tuning the pretrained Cellpose Cyto2 model under an extremely low annotation budget of only two images. Despite its conceptual simplicity and deterministic nature, CGMD achieves competitive performance, highlighting the value of principled subset selection for annotation-efficient learning. The design of CGMD is inherently model- and task-agnostic as well as domain-independent, making it applicable across a broad spectrum of datasets. Future work will focus on a systematic evaluation of CGMD with different pretrained backbones and data modalities, and on extending the comparison to more advanced subset selection and active learning strategies.

## 6. COMPLIANCE WITH ETHICAL STANDARDS

This research study was conducted using the data publicly available at https://celltrackingchallenge.net/2d-datasets/. Ethical approval was not required as confirmed by the license attached with the open access data.

## 7. ACKNOWLEDGMENTS

This research was conducted within the framework of the project “SAIL: SustAInable Lifecycle of Intelligent SocioTechnical Systems” (grant no. NW21-059B). SAIL is receiving funding from the programme “Netzwerke 2021”, an initiative of the Ministry of Culture and Science of the State of North Rhine-Westphalia, Germany. The sole responsibility for the content of this publication lies with the authors.

## Footnotes

1 celltrackingchallenge.net

